# *Vibrio aestuarianus* Clade A and Clade B isolates are associated with Pacific oyster (*Crassostrea gigas*) disease outbreaks across Ireland

**DOI:** 10.1101/2023.04.06.535062

**Authors:** Nicola M. Coyle, Ciar O’Toole, Jennifer C.L. Thomas, David Ryder, Edward J. Feil, Michelle Geary, Timothy P. Bean, Andrew Wokorac Joseph, Ava Waine, Deborah Cheslett, David W. Verner-Jeffreys

## Abstract

2.

Bacteria from the Vibrionaceae family have been implicated in mass mortalities of farmed Pacific oysters (*Crassostrea gigas*) in multiple countries, leading to substantial impairment for growth in the sector. In Ireland there has been concern that *Vibrio* have been involved in serious summer outbreaks. There is evidence that *Vibrio aestuarianus* is increasingly becoming the main pathogen of concern for the Pacific Oyster industry in Ireland. While bacteria belonging to the *Vibrio splendidus* clade are also detected frequently in mortality episodes, their role in the outbreaks of summer mortality are not well understood. To identify and characterise strains involved in these outbreaks, 43 *Vibrio* isolates were recovered from Pacific oyster summer mass mortality episodes in Ireland from 2008-2015 and these were whole genome sequenced. Among these, 25 were found to be *V. aestuarianus* (implicated in disease) and 18 *V. splendidus sensu lato* (role in disease undetermined). Two distinct clades of *V. aestuarianus* – Clade A and Clade B – were found that had previously been described as circulating within French oyster culture. The high degree of similarity between the Irish and French *V. aestuarianus* isolates points to translocation of the pathogen between Europe’s two major oyster producing countries, probably via trade in spat and other age classes. *V. splendidus* isolates were more diverse, but the data reveal a single clone of this species that has spread across oyster farms in Ireland. This underscores that *Vibrio* could be transmitted readily across oyster farms. The presence of *V. aestuarianus* Clades A and B in not only France but also Ireland adds weight to growing concern that this pathogen is spreading and impacting Pacific oyster production within Europe.

**Outcome:** Pacific oyster culture in Ireland has increasingly suffered from summer mass mortality events. Many of these mortalities in recent years have been associated with *Vibrio aestuarianus*; the role of another pathogen, *Vibrio splendidus* has, so far, remained inconclusive. Here we show that two clades of *V. aestuarianus* are circulating in Ireland, and that these are members of two clades that have previously caused extensive oyster die offs in France. Their discovery in Ireland is consistent with transport of infected oyster stock between the two countries. Although *V. splendidus-*like strains in Ireland were highly diverse, a small clonal group was detected that appears to have spread rapidly from a single source to disparate locations in Ireland. Combined, these findings highlight the appearance of a highly pathogenic *Vibrio* in Ireland, and the risk of transmission between interconnected oyster production industries in Europe.

**Data summary:** Sequences generated in this study were deposited on the NCBI. Accession number: PRJNA797364. Publicly accessed genomes are listed in Table S2.

**The authors confirm all supporting data, code and protocols have been provided within the article or through supplementary data files**.

## 5. Introduction

While the aquaculture industry has expanded rapidly in the past 50 years, oyster production has struggled to keep pace with other aquaculture products [1]. One of the significant factors constraining the development of oyster aquaculture has been infectious disease [1,2]. Pacific oysters are an important farmed species [3], with 620,000 tonnes produced on average each year worldwide between 2010 to 2019, worth an estimated US$1.29 billion a year [4]. France is the major European producer (84,760 tonnes in 2019), although there are significant industries in other European countries, including Ireland (10,460 tonnes in 2019). In France and elsewhere, there have been increased reports of disease outbreaks responsible for the depletion of oyster stocks over the last decade [3]. These present major socioeconomic consequences for the future of the oyster farming industry [5].

Episodes of abnormal mortality of Pacific oysters affecting all age classes have been described globally since the 1950s. Mortality of larvae and spat has been linked to the presence of a number of pathogenic agents including Ostreid herpes virus 1 (OsHV-1), whilst the term summer mortality syndrome has been coined to describe those events of mixed aetiology in the summer months affecting older oysters where gonad maturation is present [6]. Studies have shown that the causes of summer mortality syndrome are complex, often involving a combination of physiological and environmental stress, alongside the presence of pathogens [7], particularly bacteria belonging to the genus *Vibrio* including *V. aestuarianus* and *V. splendidus* [8].

In the summer of 2008, abnormally high mortality episodes of the Pacific oyster were reported both in France and Ireland. The losses were linked to the emergence of a new variant of OsHV-1, termed Ostreid herpes virus 1 µVar (OsHV-1 µVar) [9]. Both *V. splendidus* and *V. aestuarianus* were also detected during a number of these events [6]. Since then, mortality outbreaks have continued to spread and affect oyster farms across various parts of Europe. The frequency of detection of *V. aestuarianus* in cases of adult mortality increased significantly between 2011 and 2013, becoming the principle pathogen detected during summer mortality episodes in adult oysters in France in 2013 [10].

The Pacific oyster industry in Ireland is heavily dependent on the importation of spat which is predominantly sourced from France (Collins et al 2008). Hence, following the reports of increased detections of *V. aestuarianus* in cases of adult mortality in France, a monitoring programme and a retrospective study were instigated to determine the extent of its distribution in Ireland. In this study, we characterise and compare 43 *Vibrio* isolates recovered from diseased Irish oysters from 2008 to 2015 using whole genome sequencing.

We show, firstly, a high proportion of these oyster die-offs are associated with the presence of *V. aestuarianus* isolates from two oyster-associated *V. aestuarianus* subsp. *francensis* clades, Clade A and Clade B previously shown to be a major cause of summer mortality syndrome in France [11]. Secondly, we showcase differences in gene content diversity in these clades. Thirdly we show that *V. splendidus* strains present in Irish oysters are diverse, but a small clonal group was detected in 2009 in multiple locations.

## 6. Methods

### 6.1 Bacterial isolation and initial characterisation

43 *Vibrio* isolates obtained from oysters of varying age classes (Figure 1 and Table 1) were collected from 22 sites around the Republic of Ireland between 2008 and 2015. Isolates were recovered from either haemolymph or crushed gill tissues and characterised. In most cases, isolates were recovered from sites where there were significant ongoing mortalities taking place (Table 1). They were then stored at -80°C on cryovials using the protect™ storage system following manufacturer’s instructions.

**Table 1:** *Vibrio* strains selected for sequencing from Irish oysters

| ID | Year of extraction | Freezer Ref. | FHU bacterial ref. | Site of extraction | Sample type | Tissue sample was extracted from | Reported Mortality rate % | Age class | Species |
| --- | --- | --- | --- | --- | --- | --- | --- | --- | --- |
| 16025 | 2009 | 76/09 | S/25/09 | Loughros Beag | Ab Mort | Gill | 80-90 | 0+, 1+ | *V. aestuarianus* |
| 16028 | 2014 | 32/14 | S/17/14 AB | Woodstown | Ab Mort | Haemolymph | 60-70 | 3+ | *V. aestuarianus* |
| 16030 | 2014 | 51/14 | S/43/14 A | Kinsale | Ab Mort | Haemolymph | 90 | 2+ | *V. aestuarianus* |
| 16033 |  | 30/14 | S/16/14 A | Dungarvan | Ab Mort | Gill | 10-20 | 0+, 1+ | *V. aestuarianus* |
| 16034 | 2014 | 26/14 | S/15/14 A | Achill | Ab Mort | Gill | 10 | 0+ | *V. aestuarianus* |
| 16036 | 2015 | 47/15 | S/38/15 A | Castlemaine | Ab Mort | Haemolymph | 40 | 2+ | *V. aestuarianus* |
| 16041 | 2015 | 20/15 | S/16/15 B | Woodstown | Reposus | Gill | 1 | 1+ | *V. aestuarianus* |
| 16043 | 2015 | 58/15 | S/58/15 | Donegal Bay | Ab Mort | Haemolymph | 50-90 | 2+ | *V. aestuarianus* |
| 16044 | 2015 | 67/15 | S/65/15 B | Dungarvan | Ab Mort | Haemolymph | 15-50 | 2+ | *V. aestuarianus* |
| 16048 | 2015 | 14/15 | S/10/15 B | Dungarvan | Reposus | Haemolymph | 1 | 2+ | *V. aestuarianus* |
| 16049 | 2015 | 21/15 | S/17/15 A | Dungarvan | Reposus | Haemolymph | 1.5 | 2+ | *V. aestuarianus* |
| 16050 | 2015 | 40/15 | S/28/15 B | Woodstown | Reposus | Gill | 92 | 0+ | *V. aestuarianus* |
| 16053 | 2015 | 64/15 | S/64/15 A | Kinsale | Ab Mort | Haemolymph | 70-80 | 2+ | *V. aestuarianus* |
| 16054 | 2010 | 03/10 | S/02/10 | Poularone Creek | Ab Mort | Gill | 30-40 | 0+ | *V. aestuarianus* |
| 16056 | 2015 | 59/15 | S/59/15 | Donegal Bay | Reposus | Haemolymph | 20 | 2+ | *V. aestuarianus* |
| 16057 | 2015 | 36/15 | S/27/15 A | Dungarvan | Reposus | Haemolymph | 5.5 | 2+ | *V. aestuarianus* |
| 16058 | 2015 | 60/15 | S/60/15 A | Dungarvan | Reposus | Haemolymph | 5 | 2+ | *V. aestuarianus* |
| 16059 | 2013 | 02/13 | S/15/13 B | Kinsale | Ab Mort | Gill | 15-20 | 1+ | *V. aestuarianus* |
| 16060 | 2015 | 53/15 | S/44/15 | Donegal Bay | Reposus | Haemolymph | 2 | 2+ | *V. aestuarianus* |
| 16062 | 2015 | 96/15 | S/93/15 A | Gweedore | Ab Mort | Haemolymph | 43 | 1+ | *V. aestuarianus* |
| 16063 | 2014 | 46/14 | S/37/14 A | Kinsale | Ab Mort | Haemolymph | 90 | 1+ | *V. aestuarianus* |
| 16066 | 2013 | 25/13 | S/21/13 C | Oysterhaven | Ab Mort | Gill | 50 | 0+, 1+, 2+ | *V. aestuarianus* |
| 16067 | 2015 | 16/15 | S/11/15 B | Woodstown | Reposus | Haemolymph | 0 | 2+ | *V. aestuarianus* |
| 16070 | 2014 | 28/14 | S/14/14 B | Woodstown | Research | Haemolymph | 60-70 | 1+ | *V. aestuarianus* |
| 16071 | 2014 | 57/14 | S/52/14 A | Castlemaine | Ab Mort | Haemolymph | 30 | 2+ | *V. aestuarianus* |
| 16029 | 2009 | 68/09 | S/31/09A | Ballymacoda Bay | Ab Mort | Haemolymph | 20 | 0+ | *V. splendidus sensu stricto* |
| 16035 | 2009 | 54/09 | S/34/09 A | Clew Bay | Ab Mort | Haemolymph | 10 | 0+ | *V. splendidus sensu stricto* |
| 16037 | 2013 | 23/13 | S/31/13 A | Carlingford Lough | FH Directive | Gill | 50 | 0+ | *V. splendidus sensu stricto* |
| 16051 | 2009 | 73/09 | S/27/09 | Clew Bay | Ab Mort | Haemolymph | 75 | 0+ | *V. splendidus sensu stricto* |
| 16052 | 2013 | 07/13 | S/16/13 C | Dungloe Bay | Ab Mort | Haemolymph | 20 | 1+ | *V. splendidus sensu stricto* |
| 16065 |  | 67/09 | S/37/09 | Dungloe Bay | Ab Mort | Haemolymph | 35 | 0+ | *V. splendidus sensu stricto* |
| 16069 |  | 31/14 | S/17/14 A-A | Woodstown Strand | Ab Mort | Haemolymph | 60 | 3+ | *V. splendidus sensu stricto* |
| 16072 | 2009 | 79/09 | S/21/09 | Clew Bay | Ab Mort | Haemolymph | 3-50% | 0+ | *V. splendidus sensu stricto* |
| 16073 | 2009 | 66/09 | S/40/09 B | Valentia River | Ab Mort | Haemolymph | 45 | 0+ | *V. splendidus sensu stricto* |
| 16074 | 2008 | 34/08 | S/30/08/A | Dungarvan Harbour | Ab Mort | Haemolymph | 15 | 1+ | *V. splendidus sensu stricto* |
| 16061 | 2008 | 35/08 | S/30/08/B | Dungarvan Harbour | Ab Mort | Haemolymph | 15 | 1+ | *V. splendidus sensu stricto* |
| 16077 |  | 47/16 | S/12/16B |  |  |  |  |  | *V. splendidus sensu stricto* |
| 16078 | 2008 | 23/08 | S/028/08/B | Castlemaine Harbour | Ab Mort | Haemolymph | 85 | 1+ | *V. splendidus sensu stricto* |
| 16079 | 2015 | 63/15 | S/63/15 B | Trawenagh Bay | Ab Mort | Haemolymph | 70 | 1+ | *V. splendidus sensu stricto* |
| 16040 | 2009 | 78/09 | S/24/09B | Loughros Beag | Ab Mort | Haemolymph | 10 | 1+ | *V. splendidus sensu lato* |
| 16042 | 2013 | 28/13 | S/35/13 A | Dunmanus Bay | Ab Mort | Haemolymph | 40 | 1+ | *V. splendidus sensu lato* |
| 16075 | 2010 | 16/10 | S/42/10 | Carlingford Lough | Ab Mort | Haemolymph | 30 | 0+ | *V. splendidus sensu lato* |

**Figure 1:**
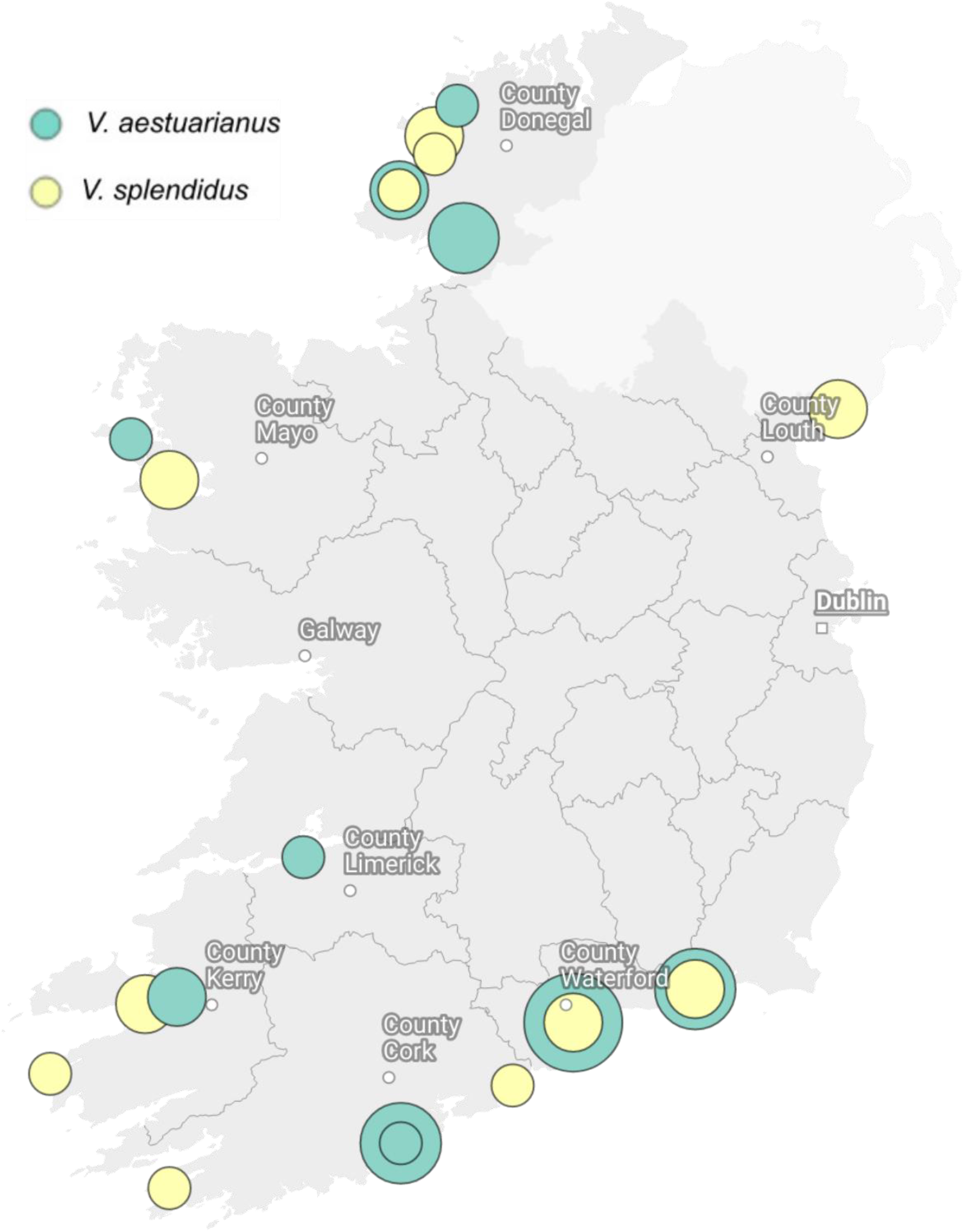
Map of 43 strains sampled across Ireland. 24 *V. splendidus* and 18 *V. aestuarianus* isolates were collected from 23 locations. Pie charts indicate the proportion of each species sequenced from each location. These nodes are weighted by the number of isolates (scale = 1 to 6).

### 6.2 DNA extraction and quantification

DNA was extracted from the isolates using the MasterPure™ Gram positive DNA extraction kit (Cat. No. MGP04100; Epicentre®). The standard protocol was modified slightly to accommodate for the isolates being Gram negative organisms. In summary, a 1 µl loopful of bacteria (previously sub-cultured onto seawater agar (SWA) was placed into a 1.5 ml Eppendorf tube containing 1 ml 0.9% saline. The solution was centrifuged at 1500 rpm, supernatant was removed and 150 µl TE buffer was added. Samples were vortexed to re-suspend the pellet. 150 µl of a premade dilution of proteinase K in Gram-positive lysis solution was added to each sample, at a concentration of 1 µl Proteinase K per 150 µl of Gram-positive lysis solution. The samples were vortexed and subsequently incubated at 65-70°C for 15 min which included vortexing every 5 min. Samples were cooled to 37°C then put on ice for 3-5 minutes, following which 175 µl of MPC Protein Precipitation Reagent was added to each sample. Samples were vortexed and centrifuged at 1500 rpm at 4°C for 10 min. The supernatant was collected (pellets discarded) and 500 µl of isopropanol was added, samples inverted 30-40 times and centrifuged again at 1500 rpm at 4°C for 10 min. The supernatant was removed, 70% ethanol was added, and samples centrifuged for a final time at 1500 rpm at 4°C for 5 min. Finally, the supernatant was removed, and samples were re-suspended in 100 µl of molecular grade water and stored at -80°C until future use. The extracted DNA was quantified using a Quantus™ fluorometer (Promega), and quality assessed using NanoDrop™ ND-1000 Spectrophotometer (Thermo). Only those samples that passed the quality check were selected for high-throughput (Illumina) sequencing.

### 6.3 Illumina Sequencing

Isolates were sequenced using an Illumina Miseq according to the standard protocols produced by the manufacturer. In brief, the DNA quantities were checked by fluorescence, diluted and prepared for sequencing with the Illumina Nextera XT library preparation kit, including optional 96-barcode adapters. Cleaned libraries were then sized-checked with an Agilent Technology 2100 Bioanalyzer using a High sensitivity DNA chip and quantified by Promega Quantus™ fluorometer using OneDNA protocol. Finally, libraries were normalised, pooled and sequenced on the Miseq with Illumina V3 600 chemistry.

### 6.4 Quality Check

Sequences were trimmed using version 0.36 of Trimmomatic, with parameters :ILLUMINACLIP:*:2:30:10 MINLEN:36 SLIDINGWINDOW:4:20 TOPHRED64 [12]. FastQC version 0.11.7 was used to check the quality of trimmed reads, and to ensure there were no significant contaminants [13].

### 6.5 Assembly and Identifying Open Reading Frames

Spades version 3.13.1 was used for assembly, with parameters: – careful –only assembler [14]. Contigs less than 500bp were removed and coverage less than 5 calculated using bwa and SAMtools v1.8 [15]. Assembled genomes were annotated using version 1.13 of Prokka, with options: – addgenes –centre XXX –mincontiglen 200 –cdsrnaolap [16]). Quality assessment of assemblies was carried out using QUAST v4.6.3 [17]. QC scores for all reads and assemblies are provided in Table S1.

### 6.6 Accessing public genomes of *V. splendidus* and *V. aestuarianus*

We obtained publicly available WGS data for *V. splendidus* and *V. aestuarianus* in order to place the isolates from Irish oysters into broader phylogenetic contexts. Thirteen *V. aestuarianus* genomes were contributed by Goudenège et al., 2015. Assembled genomes of 102 isolates previously characterised as *V. splendidus* were downloaded from the NCBI database [18]. Information on each of these isolates can be found in Table S2. All subsequent genomic analysis was done using datasets of 38 *V. aestuarianus* and 120 *V. splendidus* genomes.

### 6.7 Pangenome construction

A comprehensive pangenome of each species was constructed for both species using PIRATE [19], a toolbox for bacterial pangenomics analysis. We used Phandango version 1.3.0 [20] to visualise the distribution of gene families within each population. Core genome alignments were built using PIRATE. We used R version 3.2.3 [21] for statistical analysis and data visualisation.

### 6.8 Core genome phylogeny

Based on a 2.56 Mb core genome alignment we constructed a bootstrapped phylogenetic tree using RAxML-NG v. 0.9.0 [22] of the 38 *V. aestuarianus* isolates. For the larger *V. splendidus* dataset, we constructed a neighbour joining tree using RapidNJ [23] using a core genome of 2.97-Mb. Phylogenies were visualised using Microreact [24]. The project URLS are https://microreact.org/project/gfAsh7KuduL4xuSTDaVU5r-vibrio-aestuarianus (*V. aestuarianus*) and https://microreact.org/project/eMABqKLAPcn2QG5NEnCVor-vibrio-splendidus (*V. splendidus*). SNP distances between isolates were calculated using Disty McMatrixface 0.1.0 [25].

### 6.9 Phage prediction

We used PHAge Search Tool (PHAST) [26] to identify potential phages in Clade A isolate 12142, and Clade B isolates 01308 and 16060. Fasta assembly files were assessed using default PHAST parameters.

All Bioinformatics was carried out using resources provided by MRC-CLIMB [27].

## 7. Results

### 7.1 *Vibrio aestuarianus*: Presence of two clades in Ireland

The core genome phylogeny of *V. aestuarianus* (Figure 2) revealed that the French and Irish isolates were highly similar. The Irish isolates were resolved into the same two clades, A and B, previously reported to be circulating in French oyster culture [11]. In each clade, French isolates occur closer to the root of the tree than the Irish isolates. Strains isolated from these two countries differ by 50 SNPs on average in Clade A and 416 SNPs in Clade B.

**Figure 2:**
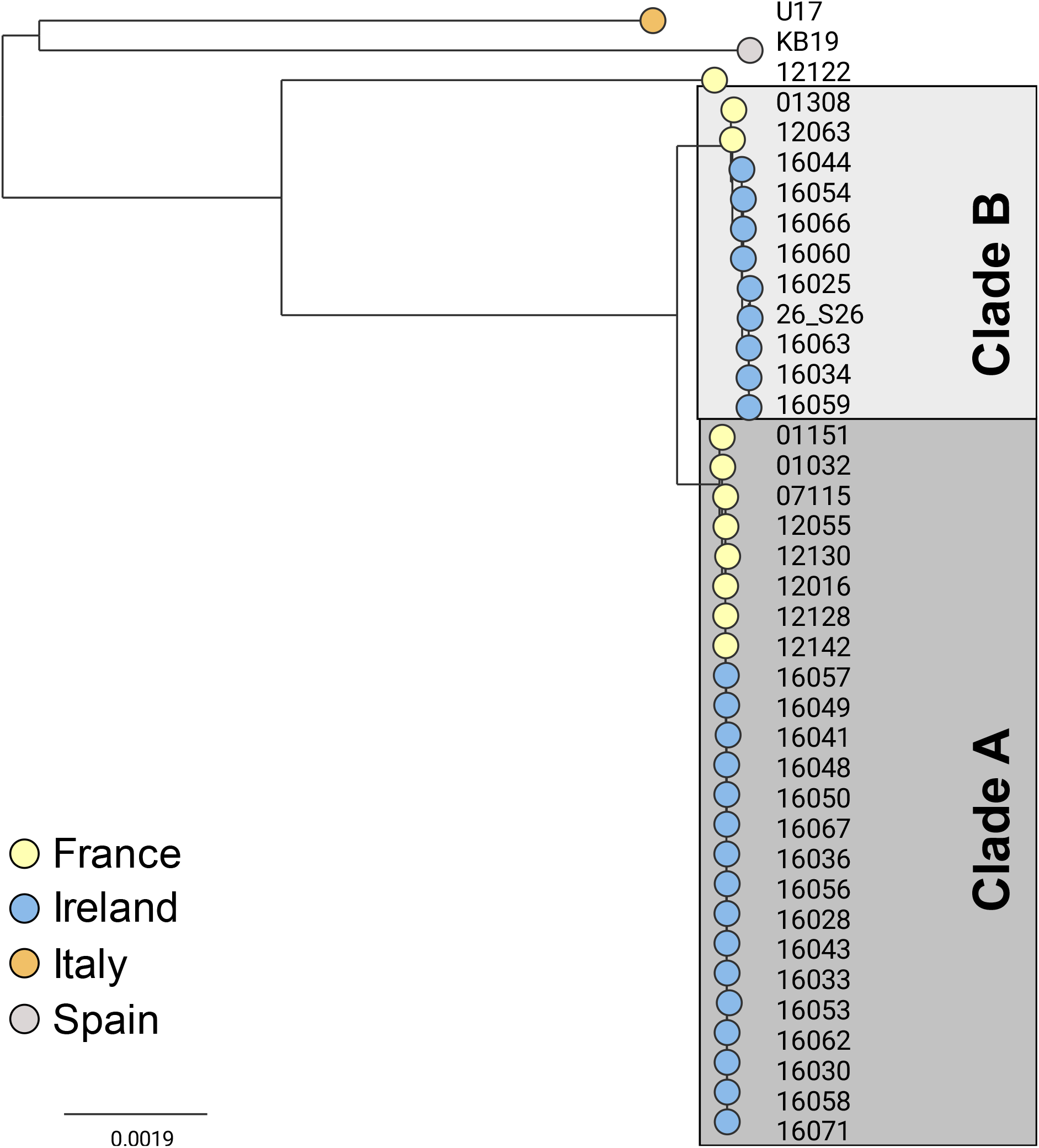
Core genome phylogeny of 38 *V. aestuarianus* isolates reveal two clades circulating in Ireland and France. Tree tips are coloured by country of isolation. Isolates recovered in Ireland fall within two previously identified clades circulating in France.

### 7.2 *Vibrio aestuarianus*: Gene content variation in each clade

The pangenome of *V. aestuarianus* consists of 5,650 gene families (Figure 3). This includes 2,746 core gene families present in at least 95% of isolates, 1,150 shared by 10%-95% isolates, and 1,754 shared by a single isolate up to 10% of isolates. Isolates 01151 and 01032 are missing many core genes due to poor quality assemblies: these were excluded from further pangenome analyses.

**Figure 3:**
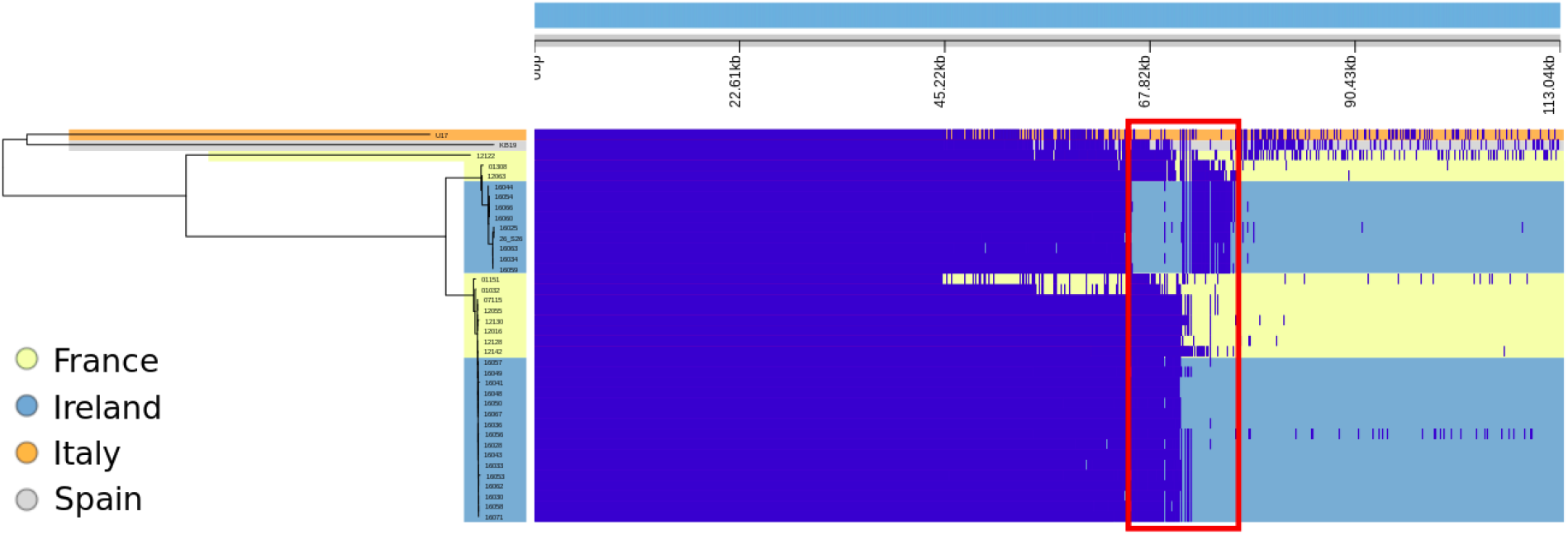
Gene presence-absence of 38 *V. aestuarianus*. Presence-absence heatmap of the pangenome of 38 *V. aestuarianus* genomes generated by Phandango [20]. Dark blue blocks indicate the presence of a gene family. Tree branches and heatmap rows are coloured by country of isolation. Indicated in a red box are multiple genes that differ between Clade A and Clade B. Isolates 01151 and 01032, French Clade B isolates, notably contain most of these genes.

A set of 215 gene families present in all Clade A isolates are absent in Clade B isolates (Figure 3). These genes are likely to have been horizontally acquired as mobile genetic elements (MGEs). To examine this, we checked the locations of these genes on the genome of the Clade A isolate 12142 and compared the GC content of these genes to the rest of the genome. The 215 genes resolved into 19 clusters, each with at least two genes (Table S3). The largest of these clusters contains 48 genes and has a GC content of 45.56%, much higher than the genome average of 42.65%. Another 13.5-Kb region with 15 genes and a GC content of 43% can be found 866-Kb away from this region on the same contig. These two large gene clusters have been identified as phages using PHAST (Table S4).

The remaining clusters of genes are distributed across 11 contigs and contain mostly hypothetical proteins (108 of 130 genes). The presence of antitoxin and phage related proteins (YafN and IntA) suggests that many of these genes may lie on other uncharacterised mobile elements or plasmids.

Clade B isolates contain 92 gene families which are not shared with Clade A, and the location of these genes was checked in Clade B isolate 16060. These are also largely hypothetical proteins (63 of 92) and are spread across 32 contigs in isolate 16060, each carrying between one and nine of these genes (Table S5). Genes related to two citrate fermentation operons which allow citrate to be used as an energy source in *V. cholerae, citCDEFXG* and *citS-oadGAB-citAB* [28], are only present in Clade B isolates. *citD-G* and *citX* are all colocalised with *citB* and *citA* (also known as *dpiA* and *dpiB*). Genes *oadA, oadB* and *oadG* are found with *citC* and copies of *citA* and *citX*. No *citS* genes were detected in this species. Genes *citA, citG*, and one copy of *citX* are also found in one Clade A genome: 12142. *vspR*, a virulence gene repressor in *Vibrio cholera* [29], is also only found in Irish Clade B genomes.

We also note that Clade B strains isolated in France harbour both sets of genes, the 92 Clade B genes, and the 215 genes that are otherwise unique to Clade A. This indicates that the Clade B strains from Ireland included in this study have experienced extensive gene loss.

### 7.3 *V. splendidus*: Widespread clonal group uncovered

To place *V. splendidus* isolates appearing in Irish oysters within the population structure of this species, we compared these 18 strains to 102 *V. splendidus* publicly available genomes. The phylogeny of *V. splendidus*-like isolates revealed a large cluster of 95 isolates with a bush-like population structure, accompanied by multiple more diverse lineages (Figure 4). Of the newly sequenced strains, 15 are found within the large cluster, while three strains lie within the broader population. Although publicly accessed genomes were all classified as *V. splendidus* species, a phylogenetic comparison with reference genomes within *V. splendidus* clade has shown that many of the more diverse isolates in this dataset are likely to have been misclassified (Figure 5). Instead, these isolates are expected to represent other species from the *V. splendidus* family. Thus, we have designated isolates 16040, 16042, and 16075 as *V. splendidus*-like isolates. A cluster of five isolates recovered from four separate locations in Ireland show high similarity within this population (Figure 6). These isolates differ by 28 SNPs on average across the core genome alignment, whereas the remaining 10 Irish isolates within the *V. Splendidus* sensu stricto cluster differ by 83, on average.

**Figure 4:**
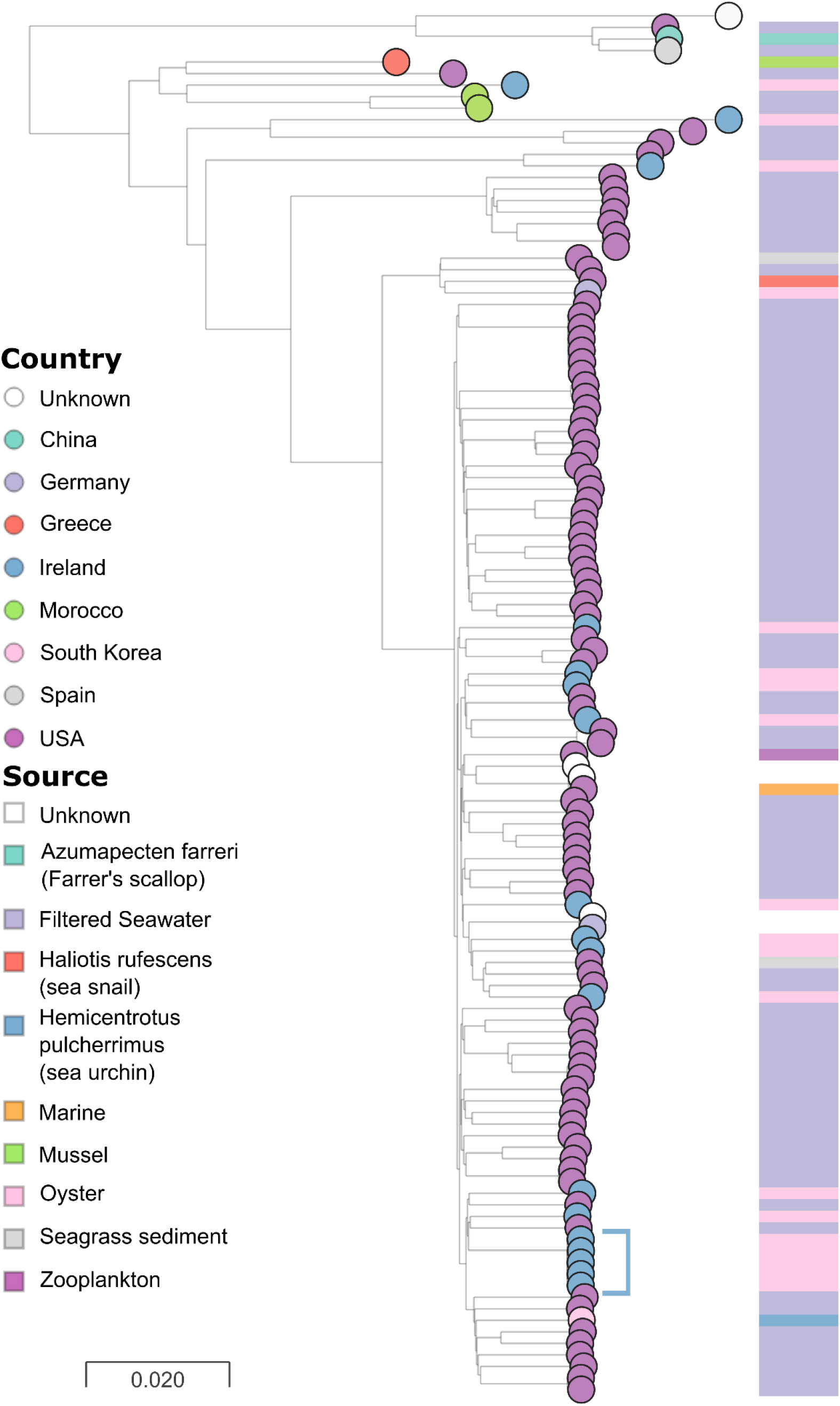
*V. splendidus* core-genome phylogeny. Neighbour-joining core-genome phylogeny of 120 *V. splendidus-like* isolates coloured by country of isolation. The tree is annotated with the source of isolation. Publicly available samples largely come from America and were sampled in seawater. The population structure of the dataset includes a large cluster of 95 genomes with a bush-like appearance. Isolates from Ireland are distributed throughout this population. However, one cluster of five highly similar isolates can be identified.

**Figure 5:**
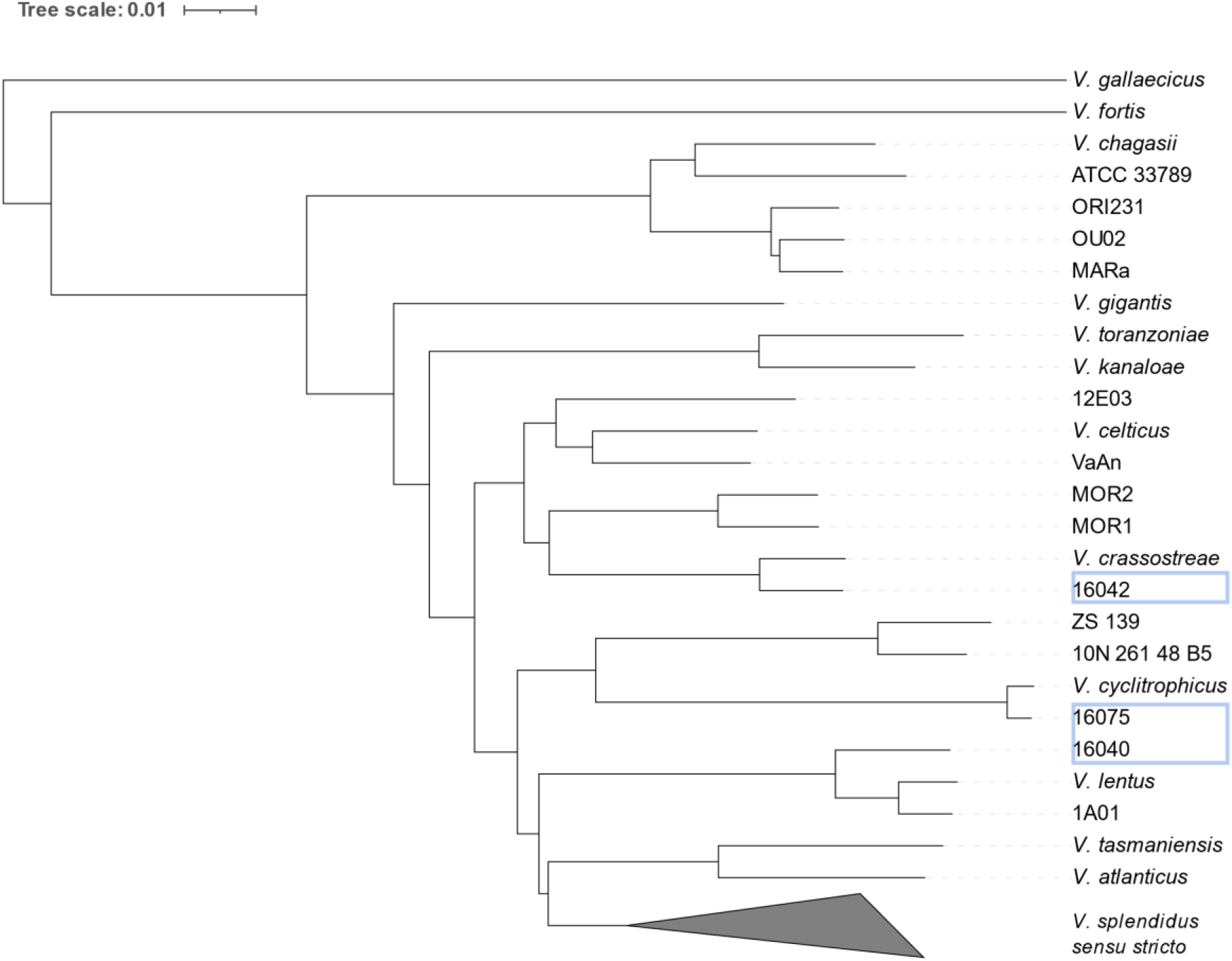
Phylogeny of *V. splendidus* species complex. The reference genomes of thirteen species belonging to the *V. splendidus* complex were combined with the 120 *V. splendidus* genomes used previously. Above is a neighbour-joining tree constructed using a core genome alignment of these genomes. *V. splendidus sensu stricto*, containing the *V. splendidus* reference strain, is collapsed and represents 107 isolates. Eleven isolates identified as *V. splendidus* species in the NCBI, that do not fall within *V. splendidus sensu stricto*, are more similar to *splendidus*-like reference genomes. Similarly, three genomes isolated in Ireland — 16075, 16040 and 16042 — are not found within *V. splendidus sensu stricto*.

**Figure 6:**
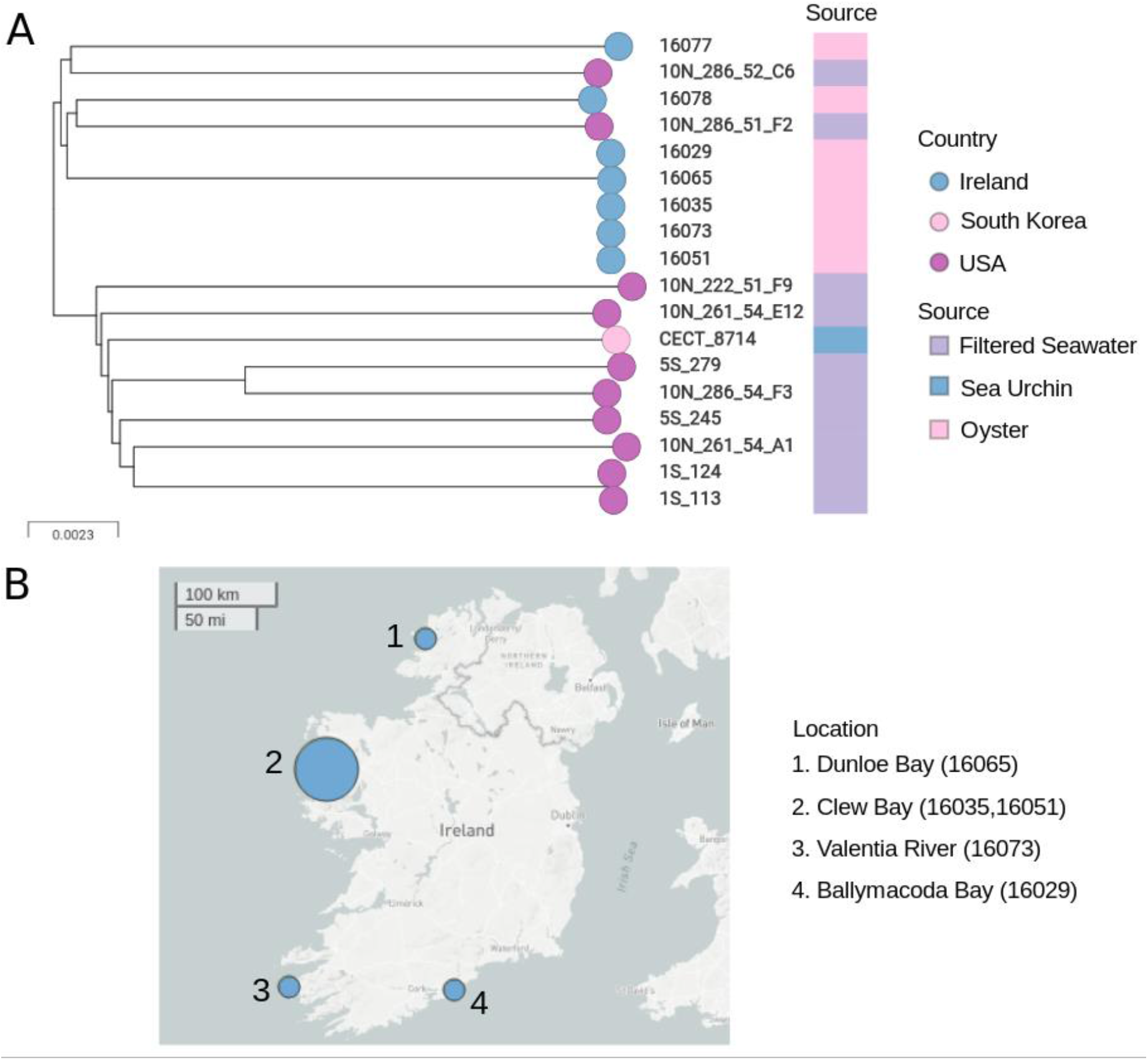
Subtree of *V. splendidus* reveals a cluster of five highly similar isolates in Ireland. A. Subtree of 19 *V. splendidus* isolates including five isolates with high similarity. Tree tips are coloured by country and annotated with the source of isolation. B. Map of the five related *V. splendidus* strains shows these isolates were not recovered in the same locations. Nodes are weighted by the number of isolates per location.

The pangenome of this species contains 18,891 gene families, with a core genome of 3,513 genes (95-100% of isolates) and 13,270 rare accessory genes (0-10% of isolates) (Figure S1). 42 gene families are unique to the five Irish clonal group isolates. These include 18 genes dispersed within a 35.6-Kb region include a trio of resistance related genes: cobalt-zinc-cadmium resistance protein, *czcA*; multidrug resistance protein, *mdtA;* and outer membrane protein *oprM*. Multiple genes related to stress response and signalling are also found in this region including *nreB* oxygen sensor histidine kinase; *cmpR* a transcriptional activator involved in CO2 stress [30]; *htpG* a chaperone protein involved in general stress responses [31]; a putative signalling protein; and *pdeB* a gene implicated in biofilm formation [32].

**Figure S1:**
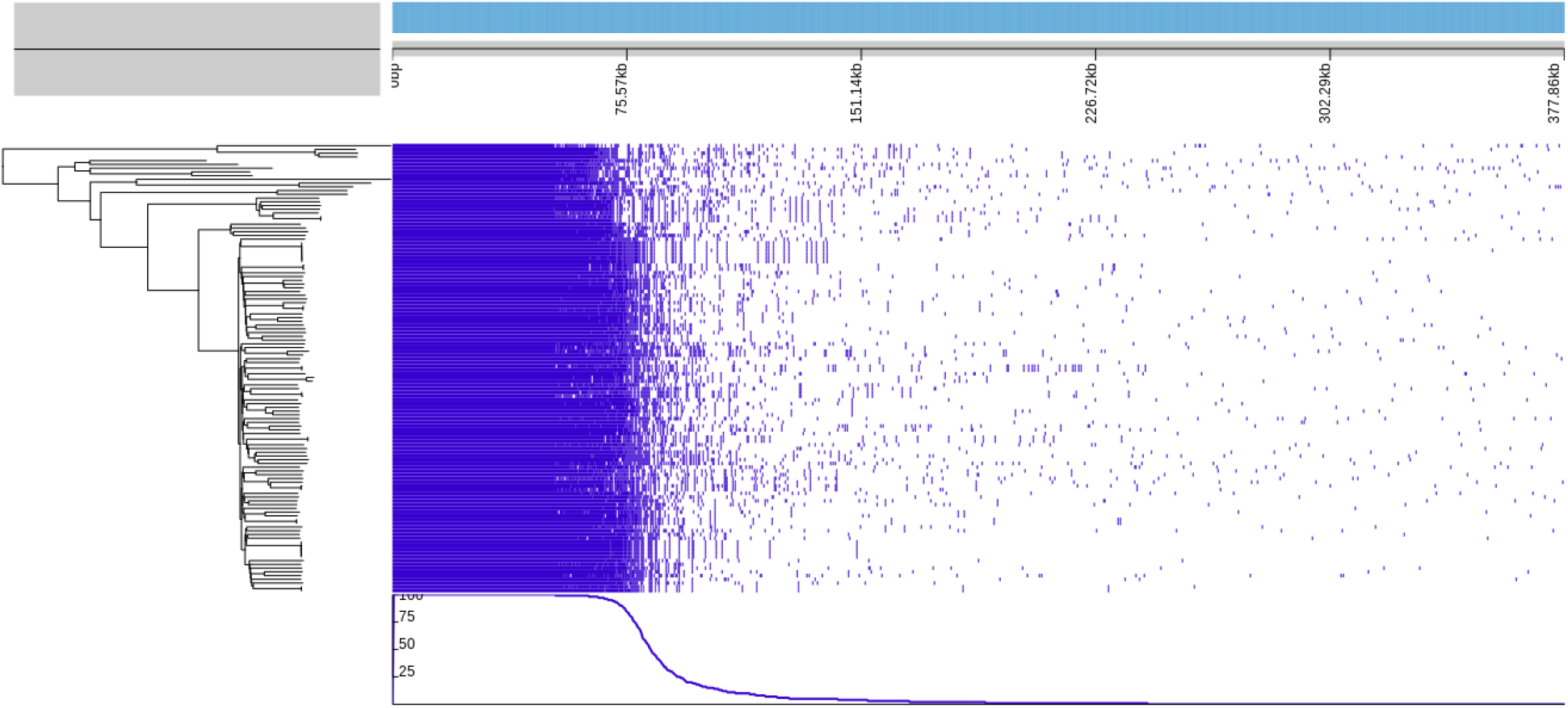
*V. splendidus* on gene presence-absence. Gene presence-absence for 120 *V. splendidus-like* isolates. Dark blue indicates the presence of a gene family.

## 9. Discussion

In Ireland, *V. aestuarianus* has been detected in oyster mortality events reported to the Marine Institute in 2001, 2003, 2006, and 2007 and more frequently in mortality events in spat from 2008 onwards, which had previously been attributed to OsHV-1 [33](D. Cheslett per. Comm.) Whilst mortality in adult oysters was only infrequently reported in Ireland prior to 2012, the frequency of reports and the detection of *V. aestuarianus* increased in line with those seen in France particularly from 2015 onwards following massive mortality events country wide in 2015 [33,34]. The trend of increased detections mirrored that in France; although the timeline of increased detections was later than that reported in France [10,34–36]. The predominant pathogen detected in cases of adult and half-grown mortality was *V. aestuarianus* whilst that in spat was OsHV-1µVar. However, other bacteria, particularly other *Vibrio* sp. have also been isolated, mainly in conjunction with OsHV-1 and *V. aestuarianus*. Here by applying whole genome sequencing we have characterised *Vibrio* strains that might play a major role in Irish oyster mortality events.

### 9.1 Two *V. aestuarianus* clades linked with oyster mortalities in both Ireland and France

Our results show that all *V. aestuarianu*s strains detected in oysters in Ireland are members of two *V. aestuarianus* subsp. *francensis* clades, A and B, which have been previously detected in France [11]. SNP analysis revealed a high level of identity between the Irish and French *V. aestuarianus* isolates, suggesting that the clades causing disease outbreaks in France are also responsible for disease outbreaks across Ireland. There is a significant trade in live oysters between France and Ireland [36,37], which has likely facilitated the movement of pathogens between rearing areas. However, broader genomic surveillance of *V. aestuarianus* associated with oyster mortalities is needed to uncover the exact distribution of each clade outside of these key *C. gigas*-producing countries.

A recent study involving the sequencing of *V. aestuarianus* strains across Europe showed that these two oyster-associated clades have now been found in multiple countries within Europe [38]. The authors found low genomic diversity within each clade and suggested that their emergence may have been the result of adaptation to oyster pathogenicity. Thus, the high genetic identity between Irish and French strains does not necessarily indicate a direct transmission chain between these two countries. While the data assessed here cannot be used to evaluate fine-scale transmission events between Ireland and France in *V. aestuarianu*s, we advocate for further whole genome sequencing efforts within and across interconnected oyster-producing countries in Europe and elsewhere to help capture the spread and evolution of these emerging infectious clades [39].

### 9.2 Evidence of gene loss in Irish Clade B strains

Our data revealed a large number of gene families that are found in French but not Irish Clade B isolates. This difference in genome content may suggest that a Clade B strain was introduced once to Ireland, and that the founder population lost or previously lacked those genes. Although some of these genes were revealed to be on phages, the mechanisms of gene loss of the remaining 152 non-consecutive gene families in these otherwise highly related strains has not been determined. It is possible that this rapid genome reduction may have conferred a selective advantage to the Irish strains [40]. Given that these Irish strains are only compared to two strains from France, more extensive sequencing of Clade B isolates across Europe and other affected regions is needed to evaluate the full diversity of the clade and determine if this gene loss is exclusive to Irish strains.

### 9.3 A single clone of *V. splendidus* highlights transmission potential

*V. splendidus* clade strains were frequently detected in Irish oyster mortalities, although the role they played in disease is uncertain. Here we showed that these isolates were mostly distinct strains within a highly diverse species complex. *V. splendidus* is a highly diverse species and opportunistic pathogen [41,42]. Given this, we would expect isolates associated with disease in Ireland to be largely unrelated, unless they happened to be isolated in the same location at one time or had recently been introduced through a common source. In 2009, a clonal group of highly similar isolates was found in multiple locations across Ireland (Figure 6). In all cases, samples were taken where mortality was occurring in recently introduced French oyster seed. Both OsHV-1 µVar and *V. splendidus* were detected, suggesting that these isolates may be linked through the source of oyster seed. While this clonal group may have proliferated across Irish waters in 2009, given that such events have not been described in this species to date, it is much more likely that it was spread to multiple farms through a common source. Indeed, at least four of these isolates were found in sites which at that time contained stock from the same hatchery in France. The occurrence of this highly related clonal group of *V. splendidus* across multiple sites in the same year signifies the presence of transmission routes available to important oyster pathogens between production facilities.

### 9.4 Perspectives

Pacific oyster summer mortality events in Ireland are shown here to be associated with two *V. aestuarianus* clades and a variety of strains within the *V. splendidus* complex. Notably, the two *V. aestuarianus* clades in Ireland have been described elsewhere in Europe, as Clade A and B [11,38]. Novel lineages where not detected which underscores the importance of these two clades in Pacific oyster summer mortalities. The occurrence of a probable transmission event of *V. splendidus* across Ireland emphasises capacity for spread of potentially pathogenic *Vibrios* within the oyster industry. Further genomic surveillance studies, which can build on this one, are needed within countries experiencing summer mortality syndrome and countries with which they frequently trade. This could lead to a fuller picture of the proliferation and evolution of this emerging pathogen and to better measures to prevent or deal with its future spread.

## 10. Author statements

### 10.1 Author contributions

NC: Formal Analysis, Investigation, Methodology, Software, Visualization, Writing – original draft, Writing – review & editing. COT: Data curation, Investigation, Writing – original draft, Writing – review & editing. JT: Investigation, Methodology, Writing – original draft, Writing – review & editing. DR: Data curation, Formal Analysis, Investigation, Methodology, Visualization, Writing – original draft, Writing – review & editing. MG: Investigation. TB: Conceptualization, Methodology. AWJ: Investigation. AW: Investigation. EF: Conceptualization, Methodology, Supervision, Writing – original draft, Writing – review & editing. DC: Conceptualization, Supervision, Writing – original draft, Writing – review & editing. DVJ: Conceptualization, Funding acquisition, Resources, Supervision, Writing – original draft, Writing – review & editing.

## 10.4 Acknowledgements

The authors would like to thank Dr Sion Bayliss and Dr Harry Thorpe for their contributions to the bioinformatics scripts used in this study. We would also like to acknowledge the contribution of the staff at the Fish Health Unit, Marine Institute, Ireland. Many thanks to Dr Georg Engelhard for his generous guidance on the manuscript. Finally, the author would like to extend gratitude to Dr Marie-Agnès Travers and colleagues at Ifremer for contributing data to this study.

